# Mutation of an insulin-sensitive *Drosophila* insulin-like receptor mutant requires methionine metabolism reprogramming to extend lifespan

**DOI:** 10.1101/2025.02.28.640731

**Authors:** Marc Tatar, Wenjing Zheng, Shweta Yadav, Rochele Yamamoto, Noelle Curtis-Joseph, Shengxi Li, Lin Wang, Andrey A Parkhitko

## Abstract

Insulin/insulin growth factor signaling is a conserved pathway that regulates lifespan across many species. Multiple mechanisms are proposed for how this altered signaling slows aging. To elaborate these causes, we recently developed a series of *Drosophila* insulin-like receptor (*dInr*) mutants with single amino acid substitutions that extend lifespan but differentially affect insulin sensitivity, growth and reproduction. Transheterozygotes of canonical *dInr* mutants (Type I) extend longevity and are insulin-resistant, small and weakly fecund. In contrast, a dominant mutation (*dInr*^353^, Type II) within the Kinase Insert Domain (KID) robustly extends longevity but is insulin-sensitive, full-sized, and highly fecund. We applied transcriptome and metabolome analyses to explore how *dInr*^353^ slows aging without insulin resistance. Type I and II mutants overlap in many pathways but also produce distinct transcriptomic profiles that include differences in innate immune and reproductive functions. In metabolomic analyses, the KID mutant *dInr*^353^ reprograms methionine metabolism in a way that phenocopies dietary methionine restriction, in contrast to canonical mutants which are characterized by upregulation of the transsulfuration pathway. Because abrogation of S-adenosylhomocysteine hydrolase blocks the longevity benefit conferred by *dInr*^353^, we conclude the methionine cycle reprogramming of Type II is sufficient to slow aging. Metabolomic analysis further revealed the Type II mutant is metabolically flexible: unlike aged wildtype, aged *dInr*^353^ adults can reroute methionine toward the transsulfuration pathway, while Type I mutant flies upregulate the trassulfuration pathway continuously from young age. Altered insulin/insulin growth factor signaling has the potential to slow aging without the complications of insulin resistance by modulating methionine cycle dynamics.

**Author Summary:** Mutations in the invertebrate insulin/IGF signaling system robustly extend lifespan. Yet these interventions often cause insulin resistance, reduce growth, and impair fertility. In contrast, a dominant gain-of-function mutation in the kinase insert domain of the Drosophila insulin/IGF receptor extends lifespan while maintaining insulin sensitivity, growth, or reproduction. Here, we demonstrate this unique mutation reprograms methionine metabolism pathway in a way that mirrors dietary methionine restriction, which is known to extend lifespan in several animals including Drosophila and mice. Genetic epistasis analysis verifies this methionine cycle inhibition is an underlying cause for how the kinase insert domain mutation slows Drosophila aging while maintaining insulin-sensitivity, growth, and fecundity.

## Introduction

Mutations of the insulin-like receptor extend lifespan in *C. elegans* and *D. melanogaster*. Slow aging in these and related manipulations of the insulin/IGF pathway is proposed to arise through mechanisms that involve proteostasis, DNA repair, translation rate, innate immunity, reproduction, growth, and mitochondrial capacity [1–3]. This multiplicity of potential causes makes it challenging to evaluate which are required to slow aging from those that merely co-occur with altered insulin/IGF signaling or are secondary consequences of the slow aging. One case of such pleiotropy involves the *Drosophila* insulin-like receptor (*dInr*) where a series of single amino acid substitutions extend lifespan but differentially affect growth and reproduction [4]. ‘Type I’ mutations produce small, long-lived adults with low fertility, and insulin weakly stimulates the phosphorylation of Akt. In contrast, a substitution in the kinase insert domain (KID) (‘Type II mutation’) robustly increases lifespan but produces full-sized adults with high fecundity, and pAkt is strongly induced by insulin [4]. This mutation of the kinase insert domain provides an opportunity to dissect how the insulin-like receptor of *Drosophila* can slow aging without the infertility, impaired growth and insulin resistance of canonical longevity mutations.

*Drosophila* has a single insulin-like receptor (*dInr*), as does *C. elegans* (*daf-2*). There are multiple insulin/IGF-like ligands for each of these receptors, encoded by seven loci in *Drosophila* [5] and by 40 loci in *C. elegans* [6, 7]. Binding and activation studies confirm *Drosophila* DILP2 and DILP5 are dInr agonists. Both stimulate the phosphorylation of Akt in a cell culture assay although along different time courses [8]. Structural analysis demonstrates three DILP5 ligands interact with dInr to produce an asymmetric *Ƭ*-shaped conformation, while a single DILP2 bound to dInr induces an asymmetric conformation of the receptor [9]. Seven insulin-like ligands in *C. elegans* are categorized as strong agonists based on how they affect dauer formation, cell division, and metabolism; while 10 INS peptides are seen as antagonists of Daf-2 function [6]. Given the pleiotropy of the *C. elegans* system, Gems and colleagues described how a range of *daf-2* mutations affected dauer, fertility, growth, and aging [10]. Alleles were categorized as Class 1 and Class 2 based on epistatic interactions with genes in the dauer formation pathway and by where their mutations fell within the Daf-2 receptor. Patel et al. [11] subsequently compared transcript profiles of *daf-2* alleles when combined with a mutation of the foxo transcription factor *daf-16*. Among alleles that robustly extended lifespan, mutants varied in how they activated Daf-16 and by which genes they induced. Patel concluded that the site of amino acid substitution within the insulin-like receptor dictates phenotypes by altering the balance of the receptor intracellular signaling [11].

Yamamoto [4] conducted an allele series analysis of the *Drosophila* insulin-like receptor by evaluating how single amino acid substitutions variously affected lifespan, growth, and reproduction. Complementation analysis was used to screen an old collection of alleles [12], where longevity candidates were subsequently regenerated by homologous recombination in a coisogenic background. Four alleles robustly extended male and female lifespan when combined as trans-heterozygotes. No alleles were viable as homozygotes while one allele, *dInr*^353^, also extended lifespan as a wildtype heterozygote (*dInr*^353^/*dInr*^wt^). All trans-heterozygotes produced small adults with reduced fecundity. In contrast, the growth and reproduction of *dInr*^353^/*dInr*^wt^ adults was equal to or greater than wildtype. Yamamoto evaluated the ability of these *dInr* genotypes to induce pAkt. Trans-heterozygotes weakly induced pAkt in an *ex vivo* insulin stimulation assay but *dInr*^353^/*dInr*^wt^ was as effective as wildtype [4].

The *dInr* longevity alleles differ by single amino acid substitutions. Alleles that extend lifespan only as trans-heterozygotes include a substitution in the extracellular Fibronectin III domain (*dInr*^E19^), a substitution in the intracellular tyrosine autoactivation loop (*dInr*^74^) and a substitution in the kinase domain C-terminal lobe (*dInr*^211^). We categorize these *Drosophila* insulin-resistant, growth-retarding, recessive longevity alleles as ‘Type I’. In contrast, the allele *dInr*^353^ is a substitution (Arg1466Cys) in the Kinase Insert Domain (KID), an unstructured peptide sequence of 27 residues that links the N- and C-lobes of the intracellular kinase domain. We categorize this *Drosophila* insulin-sensitive, growth-promoting longevity allele as ‘Type II’. Kinase insert domains occur in all tyrosine kinase receptors where they vary in length and function [13]. Unlike typical tyrosine kinase receptors, the KIDs of insulin-like receptors are short, and their functions are largely unknown. *dInr*^353^ in *Drosophila* represents the first case of non-pathogenic lesion in a KID of insulin-like receptors: it is a dominant allele that extends lifespan without insulin-resistance, reduced growth or impaired fecundity. Here we describe how *dInr* Type I and Type II alleles differentially affect the adult transcriptome and metabolome to understand the potentially unique mechanism by which *dInr*^353^ slows aging without insulin resistance.

## Results

### Transcriptional profiles of Type I dInr^E19^/dInr^74^ and Type II dInr^353^/dInr^wt^

We conducted RNAseq from somatic tissue of 10-day-old mated females of *dInr*^E19^/*dInr*^74^ (Type I), *dInr*^353^/*dInr*^wt^ (Type II), and coisogenic wildtype (*dInr*^wt/wt^ line ‘29B’). 862 transcripts were differentially expressed among the three genotypes (**Fig 1A**) (**Sup. Table 1**; iDEP http://bioinformatics.sdstate.edu/idep96/). 191 differentially expressed genes relative to wildtype were common to *dInr*^E19^/*dInr*^74^ and *dInr*^353^/*dInr*^wt^. Relative to wildtype, adult *dInr*^E19^/*dInr*^74^ uniquely altered 439 genes, while 233 were uniquely expressed by *dInr*^353^/*dInr*^wt^. The first principal components axis distinguished among all these genotypes, while the two longevity genotypes were similar but distinct from wildtype on principal component axis 2 (**Fig 1B**).

**Figure 1.**
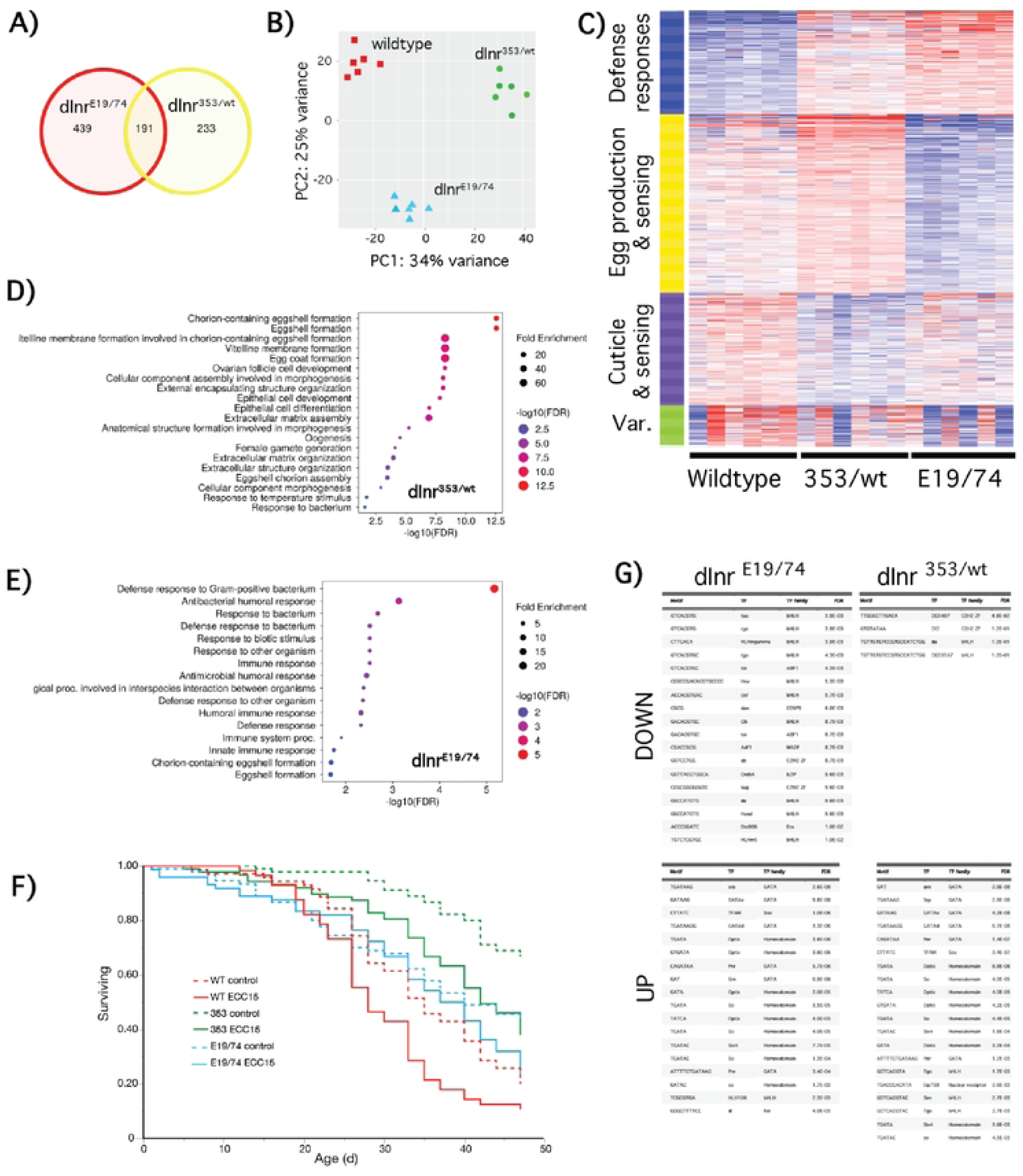
*Transcriptional profiles of Type I dInr^E19^/dInr^74^ and Type II dInr*^353^/*dInr*^wt^. (**A**) Venn diagrams of transcripts significantly altered in *Type I dInr^E19^/dInr^74^ and Type II dInr*^353^/*dInr*^wt^ genotypes. (**B**) Principal component analysis of transcriptional profiling in *Type I dInr^E19^/dInr^74^ and Type II dInr*^353^/*dInr*^wt^ genotypes. (**C**) Heatmap of transcripts significantly altered in *Type I dInr^E19^/dInr^74^ and Type II dInr*^353^/*dInr*^wt^ genotypes. (**D**) Gene set enrichment analysis of transcripts significantly altered in *Type II dInr*^353^/*dInr*^wt^ genotype. (**E**) Gene set enrichment analysis of transcripts significantly altered in *Type I dInr^E19^/dInr^74^* genotype. (**F**) Survival of *dInr*^E19^/*dInr*^74^ (Type I), *dInr*^353^/*dInr*^wt^ (Type II), and coisogenic wildtype (*dInr*^wt/wt^ line ‘29B’) infected with *ECC15* bacteria. (**G**) Down- and upregulated transcription factor binding sites in *Type I dInr^E19^/dInr^74^ and Type II dInr*^353^/*dInr*^wt^ genotypes.

K-means clustering and gene set enrichment describe how these genotypes differ based on function (**Fig 1C-E and Sup. Table 2**). Type I *dInr*^E19^/*dInr*^74^ elevates defense response genes and reduces transcripts for egg production, consistent with their low fecundity [4]. In contrast, defense response is weakly amplified in *dInr*^353^/*dInr*^wt^, while this genotype increases transcripts associated with somatic activity to support egg production, consistent with their high fecundity. To evaluate the impact of altered defense gene expression, we orally infected adults with *ECC15* bacteria. Type I *dInr*^E19^/*dInr*^74^ adults resisted infection and survived as well as uninfected controls. In contrast, *dInr*^353^/*dInr*^wt^ suffered infection-induced mortality to the same extent as infected wildtype adults (**Fig 1F**). Thus, altered innate immunity is not necessary for *dInr* mutations to extend longevity.

Differences in mRNA among the genotypes arise in part from how *dInr* regulates transcription factors. We therefore used iDEP to identify transcription factor binding sites (**Fig 1G**) that are enriched in the mutants relative to wildtype. Four TF motifs were under-represented in *dInr*^353^/*dInr*^wt^ including *daughterless* (*da*), which was the only site under-represented in *dInr*^E19^/*dInr*^74^. The *dInr* mutants shared many over-represented TF binding motifs relative to wildtype with the exception *tango*, which was under-represented in *dInr*^E19^/*dInr*^74^ yet over-represented in *dInr*^353^/*dInr*^wt^, and four sites that were unique to either *dInr*^E19^/*dInr*^74^ (*HLH106*, *dl*) or *dInr*^353^/*dInr*^wt^ (*Eip75B*, *sim*). Notable for their absence were binding sites for the transcription factors *foxo* and *Aop (Anterior open)*, which are recognized to function downstream of insulin signaling in *Drosophila* to control aging [14, 15].

Both *dInr* Types altered the mRNA of genes previously associated with extended *Drosophila* lifespan. Up-regulated longevity genes in both *dInr*^353^/*dInr*^wt^ and *dInr*^E19^/*dInr*^74^ include *tequilla*, *Sodh-1*, and *Fbp1* [16–18]. Shared down-regulated mRNA include *ImpL2*, which binds insulin ligands and extends lifespan when over-expressed [19–21], and *la costa* (*lcs*), which is suppressed by AOP when the activity of this transcription factor contributes to extended lifespan [14, 22].

Adults of *dInr*^E19^/*dInr*^74^ depress *dawdle* (*daw*) and adipokinetic hormone (*Akh*). *Dawdle* is targeted by FOXO when reduced insulin signaling extends lifespan [15]. Adipokinetic hormone has glucagon-like functions that enhance lifespan by modulating water homeostasis and metabolism [8, 15, 23]. Glycine N-methyltransferace (*gnmt*) was induced in *dInr*^E19^/*dInr*^74^ but not in Type II *dInr*^353^/*dInr*^wt^. GNMT uses S-adenosyl methionine as a cofactor to methylate glycine into sarcosine [24]. Notably, over-expression of *gnmt* extends lifespan of wildtype adults and the gene is required for reduced insulin activity (via a dominant negative expression of *dInr*) to slow aging [25, 26]. We likewise find *dInr*^E19^/*dInr*^74^ increases insulin-like peptides *ilp3* and *ilp5* mRNA, indicating these adults are hyper-insulinemic.

Few transcripts with recognized impact on aging change exclusively in *dInr*^353^/*dInr*^wt^. Rather, many DEGs of *dInr*^353^/*dInr*^wt^ are associated with egg production, however the transcription factor daughterless (*da*) is reduced only in *dInr*^353^/*dInr*^wt^. *dInr*^353^/*dInr*^wt^ also upregulates several Tot secreted stress-response proteins (TotA, TotC and TotX), while it downregulates the innate immune proteins DptA, drosomycin, and edin.

Overall, while Type I and Type II *dInR* mutants have some common downstream transcriptional impacts, the *dInr*^353^/*dInr*^wt^ mutant also drives a program distinct from insulin-resistant Type I mutants.

### Metabolomic profiles of dInr^E19^/dInr^74^ and dInr^353^/dInr^wt^

To study the metabolic consequences of altered insulin receptor function, we performed targeted steady-state metabolite profiling of Type I insulin-resistant *dInr*^E19^/*dInr*^74^ and Type II insulin-sensitive *dInr*^353^/*dInr*^wt^ at two different ages. Somatic tissues of 15-day-old and 30-day-old-females were analyzed with six biological replicates (each of 10 females without ovary) for both longevity genotypes relative to wildtype. To analyze the effect of genotype on the steady-state metabolite profile, we first analyzed the results of 15-day-old females. We found that each genotype can be distinguished with principal component analysis (PCA) (**Sup Fig 2A**) based on the 170 detected metabolites within the target set (361 metabolites). Relative to wildtype, 26 metabolites were uniquely altered in Type II *dInr*^353^/*dInr*^wt^ and 38 were differentially abundant just in Type I *dInr*^E19^/*dInr*^74^ (FDR < 0.05). Seventeen metabolites were similarly affected in the same direction in both genotypes (**Fig 2A,B and Sup. Table 3**).

**Figure 2.**
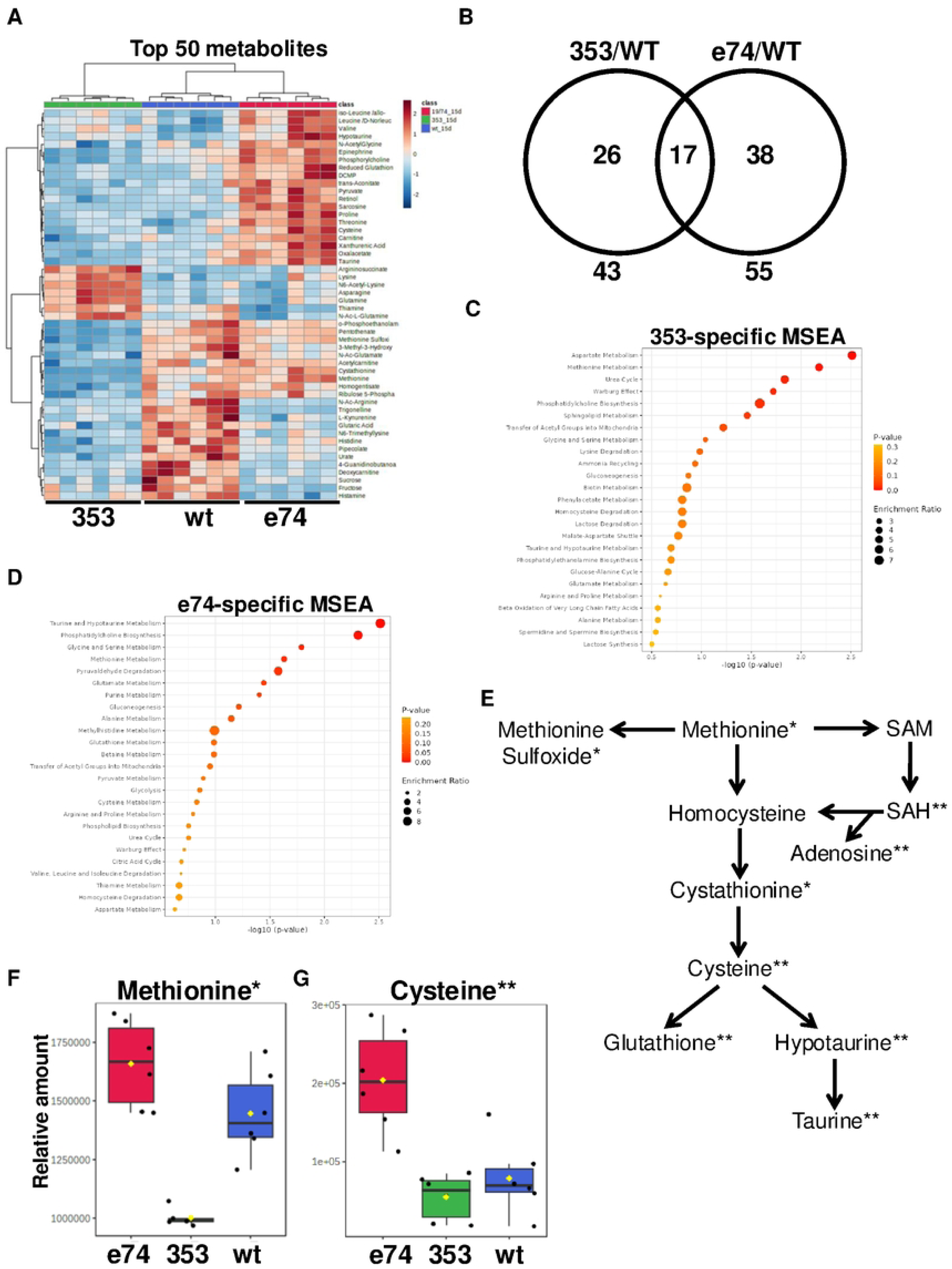
Metabolomic profiles of dInr^E19^/dInr^74^ and dInr^353^/dInr^wt^. (**A**) Heatmaps of the top 50 most altered metabolites in *Type I dInr^E19^/dInr^74^ and Type II dInr*^353^/*dInr*^wt^ genotypes. (**B**) Venn diagrams of metabolites significantly altered in *Type I dInr^E19^/dInr^74^ and Type II dInr*^353^/*dInr*^wt^ genotypes. (**C**) Metabolite set enrichment analysis (MSEA) of metabolites significantly altered in *Type II dInr*^353^/*dInr*^wt^ genotype. (**D**) Metabolite set enrichment analysis (MSEA) of metabolites significantly altered in *Type I dInr^E19^/dInr^74^* genotype. (**E**) Scheme of methionine metabolism. Significantly altered metabolites are marked with (*) in *Type II dInr*^353^/*dInr*^wt^ genotype and *Type I dInr^E19^/dInr^74^* genotype (**). (**F**) Relative levels of methionine in *dInr*^E19^/*dInr*^74^ (Type I), *dInr*^353^/*dInr*^wt^ (Type II), and coisogenic wildtype (*dInr*^wt/wt^ line ‘29B’) flies. (**G**) Relative levels of cysteine in *dInr*^E19^/*dInr*^74^ (Type I), *dInr*^353^/*dInr*^wt^ (Type II), and coisogenic wildtype (*dInr*^wt/wt^ line ‘29B’) flies.

Metabolite Set Enrichment Analysis (MSEA) distinguishes pathways affected by each mutant. Type II *dInr*^353^/*dInr*^wt^ impacts ‘aspartate metabolism’ and ‘methionine metabolism’. Insulin-resistant *dInr*^E19^/*dInr*^74^ mutants affect ‘taurine and hypotaurine metabolism’, and ‘phosphatidylcholine biosynthesis’ (**Fig 2C,D**). Notably, these longevity genotypes intersect at features of methionine metabolism (**Fig 2E**) but they do so in strikingly different ways. Type II *dInr*^353^/*dInr*^wt^ has reduced steady-state methionine, methionine sulfoxide, and cystathionine (**Fig 2E,F**; **Sup Fig 2B**) while Type I *dInr*^E19^/*dInr*^74^ has elevated cysteine, hypotaurine, taurine, and adenosine, and reduced-glutathione (**Fig 2E,G**; **Sup Fig 2D**). Sarcosine is also strongly enriched (7-fold) in Type I *dInr*^E19^/*dInr*^74^ yet less abundant in Type II *dInr*^353^/*dInr*^wt^ adults. Sarcosine is notable because this metabolite is generated by GNMT, where *gnmt* mRNA is 3.3-fold greater in *dInr*^E19^/*dInr*^74^. Previous work has established a role for *gnmt* in how impaired insulin signaling extends lifespan [25, 26]. Overall, Type I and Type II *dInr* mutants present different patterns of reprogrammed methionine metabolism.

### Age dynamics of dInr^E19^/dInr^74^ and dInr^353^/dInr^wt^ metabolic profiles

Metabolic profiles intrinsically change with age [27], including that of *Drosophila* methionine metabolism [28, 29]. We therefore analyzed how aging affects metabolic changes in *dInr*^wt^/*dInr*^wt^, *dInr^E19^/dInr^74^, and dInr*^353^/*dInr*^wt^ females using the metabolomics dataset of **Fig 2**. We first normalized levels in young flies to 100% in each genotype and analyzed how aging affects metabolite levels within each genotype (**Fig 3B**). Relative to young adults, methionine, sarcosine, and cystathionine decreased with age in wildtype *dInr*^wt^/*dInr*^wt^ (**Fig 3C,D,E**), as seen previously with wildtype *OregonR*, *yw*, and *w^1118^* flies [28, 30]. Methionine also decreased with age in *dInr*^E19^/*dInr*^74^ as did sarcosine (**Fig 3C,D**), which we previously found was substantially elevated in young Type I mutants. In contrast, in *dInr*^353^/*dInr*^wt^ adults the steady state level methionine, which was low in young adults, remained low in aged adults (**Fig 3C**). SAM increased with age in wildtype flies but remained effectively constant across age in *dInr*^E19^/*dInr*^74^ and *dInr*^353^/*dInr*^wt^ **(Fig 3F**). The level of methionine sulfoxide was relatively low in young *dInr*^353^/*dInr*^wt^ and significantly increased with age but remained constant across age in *dInr*^E19^/*dInr*^74^ and wildtype adults (**Fig 3G**).

**Figure 3.**
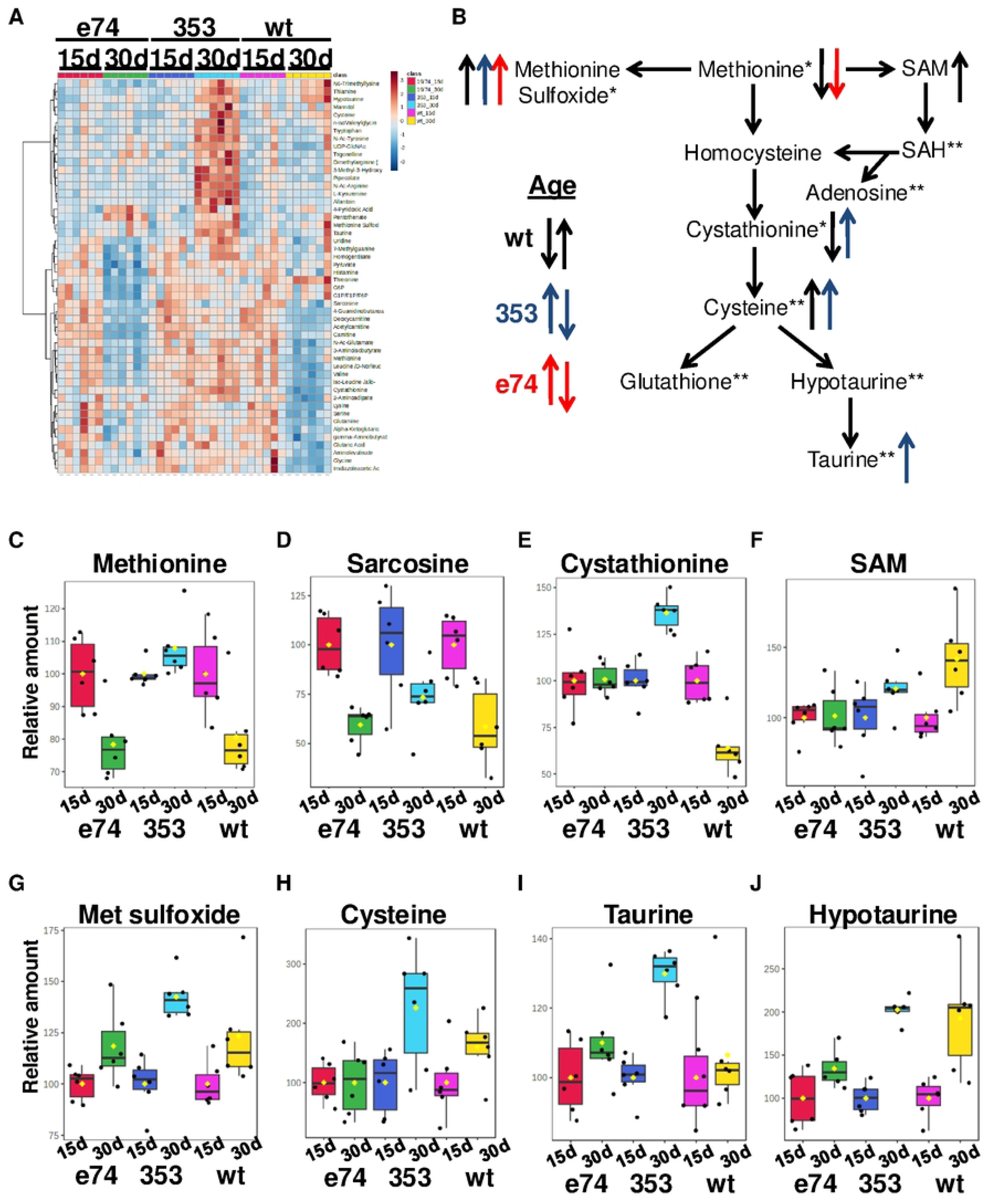
Age dynamics of dInr^E19^/dInr^74^ and dInr^353^/dInr^wt^ metabolic profiles. (**A**) Heatmaps of the top 50 most altered metabolites with age in wildtype (*dInr*^wt/wt^ line ‘29B’), *Type I dInr^E19^/dInr^74^, and Type II dInr*^353^/*dInr*^wt^ genotypes. The levels of each metabolite at a young age are normalized to 100% to reflect the age-related changes in each genotype. (**B**) Scheme of methionine metabolism. Significantly altered metabolites with age are marked with arrows. Relative changes with age of (**C**) methionine, (**D**) sarcosine, (**E**) cystathionine, (**F**) SAM, (**G**) methionine sulfoxide, (**H**) cysteine, (**I**) taurine, and (**J**) hypotaurine.

There are notable differences among the genotypes in how metabolites of the transsulfuration pathway change with age. Aged wildtype flies have low levels of cystathionine but elevate its derivative, hypotaurine (**Fig 3E,H,I,J**). The levels of transsulfuration metabolites were initially elevated in young *dInr*^E19^/*dInr*^74^ and remained high in aged adults (**Fig 3E,H,I,J)**. In contrast to the Type I genotype, all assayed transsulfuration metabolites strongly increased with age in Type II *dInr*^353^/*dInr*^wt^, including taurine (**Fig 3E,H,I,J**).

The *dInr* genotypes appear to have different age-associated changes in methionine metabolism. As a result, *dInr*^E19^/*dInr*^74^ enhances the transsulfuration pathway as young adults and maintains this high level with age. Young insulin-sensitive *dInr*^353^/*dInr*^wt^ adults produce few transsulfuration metabolites but increase these peptides as they age.

### Methionine cycle dynamics of dInr^E19^/dInr^74^ and dInr^353^/dInr^wt^

Tracking ^13^C can estimate the transition rates across steps in the methionine cycle and quantify the relative flow of methionine derivatives toward nucleotides, polyamines, and the transsulfuration pathway [31, 32]. In this approach, carbon isotope-labeled nutrients are fed to cells or organisms, and the source of carbons within specific metabolites is estimated from their relative number of labeled and unlabeled carbons [33–35].

We fed young *dInr* adults with stable isotope-labeled methionine (^13^C5-methionine, 5 labeled carbons: m+5) and measured how these carbons were incorporated into the methionine that is regenerated from homocysteine (seen as ^13^C4-methionine, 4 labeled carbons: m+4), as previously described [28, 30]. The methionine m+4/m+5 ratio is higher in *dInr*^E19^/*dInr*^74^ than in wildtype (**Fig 4B**), indicating Type II *dInr* cells have elevated methionine cycle activity. Such activity can account for their elevated steady-state sarcosine and of transsulfuration pathway products. These data imply that somatic cells of Type I mutants process an abundant amount of methionine. In contrast, the ratio of m+4/m+5 methionine is reduced in *dInr*^353^/*dInr*^wt^ relative to wildtype (**Fig 4B**), indicating these cells have lower activity across their central methionine cycle, produce less Hcy and thus have less recycled methionine. An alternative (nonexclusive) explanation is that methionine in Type II mutants is rerouted towards the TSP or the salvage pathway.

**Figure 4.**
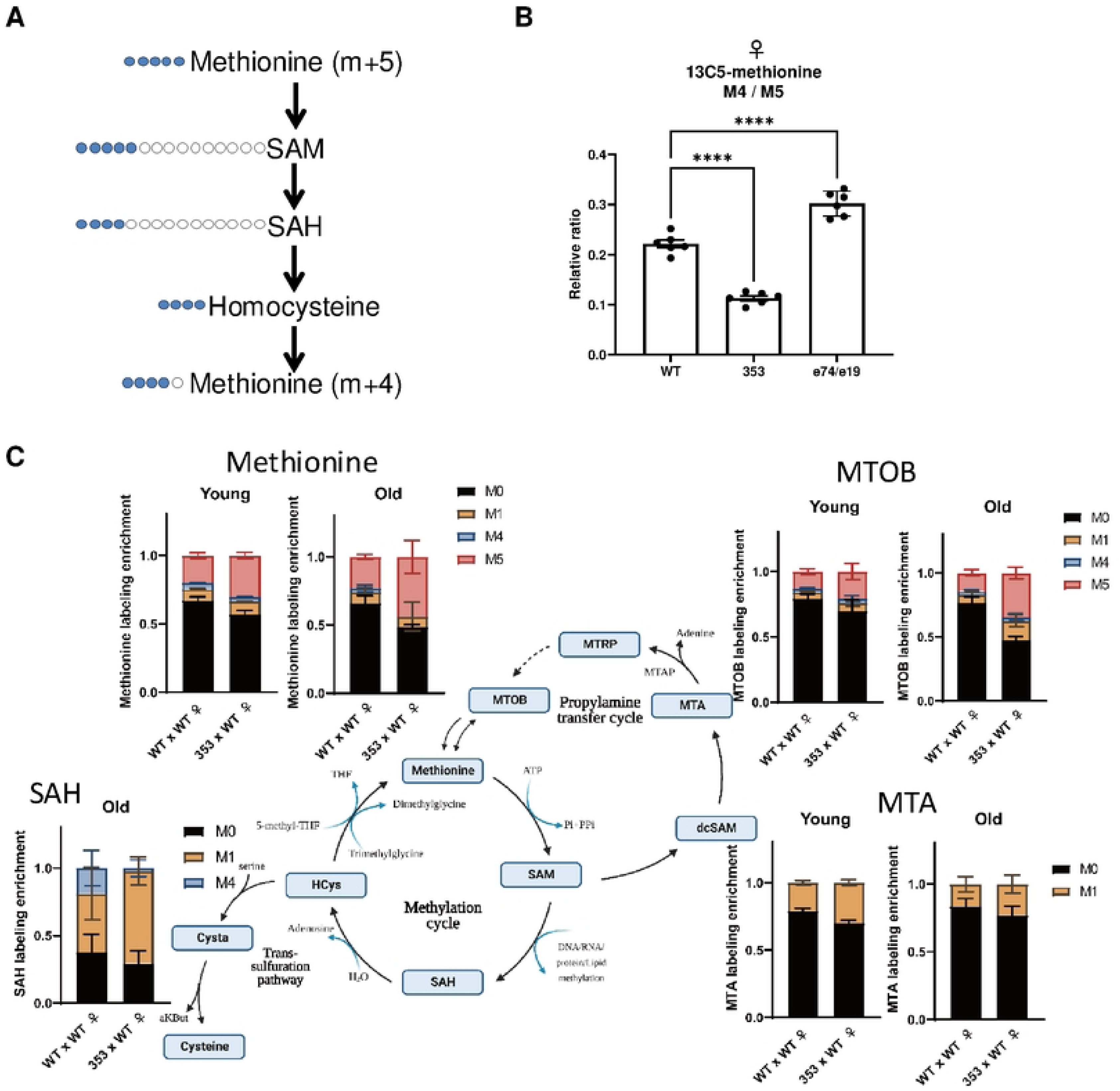
Methionine cycle flux of dInr^E19^/dInr^74^ and dInr^353^/dInr^wt^. (**A**) Scheme of methionine labeling. Blue color marks the labeled carbons. (**B**) The m+4/m+5 ratio between methionine (m+4) produced from the methionine cycle and methionine (m+5) received from the chemically-defined food in female wildtype (*dInr*^wt/wt^ line ‘29B’), *Type I dInr^E19^/dInr^74^, and Type II dInr*^353^/*dInr*^wt^ genotypes. (**C**) The distribution of labeled methionine in young and old female wildtype (*dInr*^wt/wt^ line ‘29B’), *Type I dInr^E19^/dInr^74^, and Type II dInr*^353^/*dInr*^wt^ genotypes fed with 1 mM of labeled ^13^C5-methionine tracer in chemically-defined food (lacking endogenous methionine) for 5 days. M0-M12 mark the number of labeled carbons. Labeling enrichment is the proportion of a particular labeled metabolite form among all measured isoforms. Note that although multiple isoforms of the same metabolite with different numbers of labeled carbons were measured, many of them are not present in *Drosophila* cells, and they are not displayed on the graph

It seems unlikely that methionine in the Type II is rerouted to the transsulfuration pathway because we do not observe elevated taurine or glutathione in our steady state metabolomic analysis. Because metabolites of the salvage/polyamine pathway were not well represented in the panel of the steady state analysis, to evaluate the potential rerouting of methionine into this branch we traced labeled carbons from methionine into intermediaries of the methionine cycle and salvage pathway in young and aged flies (*dInr*^wt^/*dInr*^wt^ and *dInr*^353^/*dInr*^wt^ genotypes). Dietary methionine is first converted to S-adenosyl-L-methionine (SAM), which can be decarboxylated by AdoMet decarboxylase into decarboxylated SAM (dcSAM), which serves as an aminopropyl group donor. Spermidine synthase and spermine synthase use this aminopropyl donor to covert putrescine to spermidine and spermine, while dcSAM is converted to MTA. MTA then undergoes six enzymatic steps to regenerate methionine (generating 4-Methylthio-2-oxobutanoic acid (MTOB) as one intermediary). In *dInr*^353^/*dInr*^wt^ adults we see that both MTA and MTOB are labeled with one carbon (m+1) derived from methionine, which can arise from m+4 or m+5 labeled methionine (Fig 4C). Furthermore, we see a significantly elevated fraction of m+1 labeled MTA (in young and old flies) and of m+1 labeled MTOB (in old flies), suggesting this genotype may indeed reroute methionine towards polyamine production.

Although of interest, our technical platform did not permit us to detect differential labeling of SAM and cystathionine. We detected labeled SAH only in old flies, perhaps because SAH is intrinsically elevated with age [28]. SAH generated directly from m+5 or m+4 labeled methionine will be labeled as m+4 SAH. SAH generated from methionine that passed through the salvage pathway will be labeled as m+1 SAH (via m+1 methionine). Notably, we found an elevated fraction of m+1 SAH in *dInr*^353^/*dInr*^wt^ adults and almost no fraction of m+4 SAH. These results imply that carbons from methionine in adults of *dInr*^353^/*dInr*^wt^ transits through the salvage pathway first and then enter the methionine cycle, consistent with the observed m+1 labeling of salvage pathway intermediaries (MTA and MTOB).

Overall, our data reveal two unique features of methionine metabolism in *dInr*^353^/*dInr*^wt^ adults: their central methionine cycle has reduced activity, as occurs in dietary methionine restriction, and they activate their methionine salvage pathway, which is expected to generate polyamines.

### Interactions between dInr^353^/dInr^wt^ and genes of the methionine cycle

Previous work established that reduced insulin signaling confers longevity in part because it induces *gnmt*, which itself is required for *dInr* genotypes to extend lifespan [25, 26]. Consistent with those reports, Type I *dInr*^E19^/*dInr*^74^ increases *gnmt* mRNA and sarcosine, the product of GNMT enzymatic activity. In contrast, these features were not seen in Type II *dInr*^353^/*dInr*^wt^ (**Sup. Table 1**), and correspondingly, knockdown of *gnmt* mRNA in *dInr*^353^/*dInr*^wt^ did not inhibit the exceptional longevity of the Type II mutant (**Fig 5A**). We therefore tested other components of the methionine cycle. As noted, S-adenosyl-methionine (SAM) was reduced in *dInr*^353^/*dInr*^wt^, and SAM has been recently shown in mammalian cells to modulate TOR activity by disrupting the SAMTOR-GATOR1 complex [36]. Reduced SAM promotes the association of SAMTOR with GATOR1 and inhibits TORC1. Thus, reduced SAM may promote longevity by activating SAMTOR. To test this potential interaction in *Drosophila* we depleted *samtor* in wildtype and *dInr*^353^/*wt* adults. This, however, did not diminish the extended longevity of *dInr*^353^/*dInr*^wt^ females (**Fig 5B**) while in males, *samtor*-RNAi alone repressed survival in wildtype flies (**Fig 5E**). To further test if *dInr*^353^ interacts with TOR, we overexpressed activated S6K (UAS-S6K-TE) in *dInr*^353^ and wildtype adults. This manipulation depressed survival in all genotypes but did not eliminate the relative longevity benefit of *dInr*^353^ (**Fig 5C,F**). Further downstream, S-adenosyl-homocysteine (SAH) is generated when SAM contributes a methyl-group to substrates [37]. SAH itself is an inhibitor of SAM-dependent methylation and is therefore robustly regulated by the activity of AHCY [38]. Here we find that depletion of *Ahcy13* in the *dInr*^353^/*wt* background eliminates the survival advantage of the *dInr* mutant relative to similarly treated wildtype: *dInr*^353^/*wt* requires *Ahcy* to extend lifespan in females and males. Because lifespan is restored to wildtype by blocking Ahcy13 (SAHH), we conclude the extended lifespan of the Type II *dInr*^353^/*dInr*^wt^ mutant requires reduced flux of SAH: Type II *dInr*^353^/*dInr*^wt^ slows aging by reprogramming the methionine cycle.

**Figure 5.**
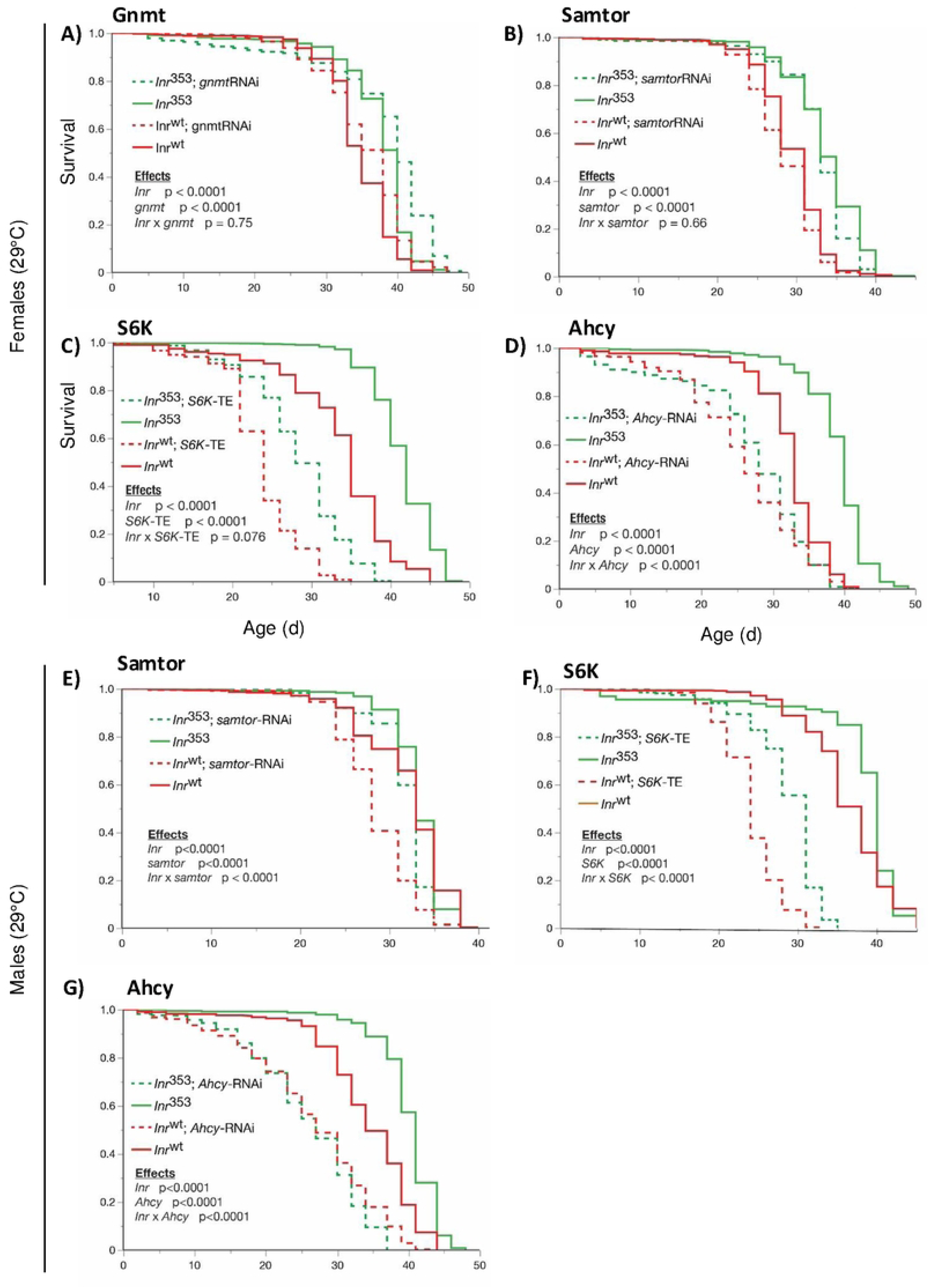
Genetic interaction between the longevity mutation of the dInr KID and the methionine cycle. (**A**) Survival of female wildtype (*dInr*^wt/wt^ line ‘29B’) and *Type II dInr*^353^/*dInr*^wt^ genotypes combined with Da-GS-Gal4 and *gnmt* RNAi fed with either EtOH or RU486. (**B**) Survival of female wildtype (*dInr*^wt/wt^ line ‘29B’) and *Type II dInr*^353^/*dInr*^wt^ genotypes combined with Da-GS-Gal4 and *samtor* RNAi fed with either EtOH or RU486. (**C**) Survival of female wildtype (*dInr*^wt/wt^ line ‘29B’) and *Type II dInr*^353^/*dInr*^wt^ genotypes combined with Da-GS-Gal4 and *S6K* transgene fed with either EtOH or RU486. (**D**) Survival of female wildtype (*dInr*^wt/wt^ line ‘29B’) and *Type II dInr*^353^/*dInr*^wt^ genotypes combined with Da-GS-Gal4 and *Ahcy13* RNAi fed with either EtOH or RU486. (**E**) Survival of male wildtype (*dInr*^wt/wt^ line ‘29B’) and *Type II dInr*^353^/*dInr*^wt^ genotypes combined with Da-GS-Gal4 and *samtor* RNAi fed with either EtOH or RU486. (**F**) Survival of male wildtype (*dInr*^wt/wt^ line ‘29B’) and *Type II dInr*^353^/*dInr*^wt^ genotypes combined with Da-GS-Gal4 and *S6K* transgene fed with either EtOH or RU486. (**G**) Survival of male wildtype (*dInr*^wt/wt^ line ‘29B’) and *Type II dInr*^353^/*dInr*^wt^ genotypes combined with Da-GS-Gal4 and *Ahcy13* RNAi fed with either EtOH or RU486.

## Discussion

### Type I and II dInr

Type I and II insulin-like receptor mutants extend lifespan, yet these genotypes differ in many ways. Typical of manipulations that reduce insulin signaling and extend lifespan, Type I *dInr* genotypes retard growth, slow development, repress fecundity, and cause insulin-resistance. They extend longevity only as transheterozygotes. Yamamoto et al. identified three alleles with these characteristics: *dInr*^E19^, *dInr*^211^, and *dInr*^74^ [4]. The *dInr*^E19^ allele was originally reported to increase lifespan and to reduce insulin receptor phosphorylation when combined with an enhancer trap insertion in the first coding exon [39], The *dInr*^E19^ allele substitutes aspartic acid for valine in the linker sequence between the L2 and FnIII-1 ectodomains. *dInr*^74^ generates a missense mutation in the conserved activation loop of the intracellular kinase domain and likely reduces the ability of the kinase to be transactivated. The *dInr*^211^ allele substitutes glycine with arginine at residue 1598 of the kinase domain C-terminal lobe. It extends lifespan when combined with *dInr*^E19^ or with *dInr*^74^, where all transheterozygotes are insulin resistant.

In contrast, the Type II *dInr* mutant of the KID has normal growth, high fecundity and robust insulin sensitivity, and it extends lifespan as a wildtype heterozygote (*dInr*^353^/*dInr*^wt^). All tyrosine kinase receptors have KIDs, which are unstructured peptide sequences that link the kinase domain N- and C-lobes [13]. KIDs often contain binding motifs for adaptor proteins although none are known in the KID of Drosophila InR, which is relatively small (27 residues), or in the KIDs of mammalian IR and IGF1R (both with 15 residues). These insulin/IGF KIDs have conserved residues at and around the start of the KID, including the arginine in *Drosophila* (R1466) that extends longevity in *dInr*^353^ when substituted for cysteine. No function to date has been identified for the KID of any insulin/IGF-like receptor. We therefore characterized transcriptional and metabolomic profiles of the Type II *dInr* relative to wildtype and the Type I mutants. We found the altered KID of dInR produces a methionine restriction-like state, and showed this mutant receptor genetically interacts with a key enzyme of the methionine cycle to slow aging.

### Transcriptional profiles of insulin-like receptor genotypes

*C. elegans* and *Drosophila* have been extensively characterized for how mutant insulin receptors (*daf-2*, *dInr*) impact mRNA profiles and enrich for daf16/foxo transcription factor binding sites [15, 40]. Working with variants of *daf-2*, Patel et al. [11] described how different alleles impact levels of mRNA. The weak allele *daf-2*(m577) modestly increases *C. elegans* lifespan and mildly affects the temperature at which animals enter dauer, while the often-studied *daf-2*(e1370) allele robustly increases lifespan, strongly affects dauer, and reduces feeding and movement [10]. Patel et al. [11] discriminated many differentially expressed genes among these longevity alleles and how these patterns depended on *daf-16*. Transcripts upregulated by *daf-2*(e1370) that were not affected in *daf-2*(m577) included small heat shock proteins, the insulin ligand *ins-22* and the nuclear hormone receptor *nhr-206*. These genes may contribute to how *daf-2*(e1370) slows aging in ways that are distinct from the action of the *daf-2*(m577) allele.

Here we discriminate among the differentially expressed genes of *Drosophila dInr* longevity mutants. Type I *dInr*^E19^/*dInr*^74^ and Type II *dInr*^353^/*dInr*^wt^ together expressed 862 different genes relative to wildtype. The Type II KID mutant *dInr*^353^/*dInr*^wt^ uniquely altered 233 transcripts that were not seen to change in the Type I. Principal components axis 1 (PC1) partitioned all the *dInr* genotypes, while the mutant genotypes were similar on PC2 but together distinct from wildtype. Type I and Type II *dInr* mutants produce common and unique transcriptional changes that may contribute to extended lifespan.

Transcripts elevated in both *dInr* genotypes include *tequila*. Previous work found a *tequila* hypomorph extended *Drosophila* lifespan, likely because it reduces neuronal insulin secretion [17]. The *dInr* mutants may elevate *tequila* to increase insulin secretion to compensate for insulin receptor resistance. Compensation may also explain why *ImpL2* is reduced in both genotypes: ImpL2 is a peptide binding protein that inhibits the biological availability of *Drosophila* insulin-like peptides [19]; limiting this inhibitor may increase insulin ligand bioavailability. Consistent with these explanations, mRNA for insulin-like ligands *ilp3* and *ilp5* is elevated in Type I, although not in Type II. Overall, Type I *dInr* adults have three features associated with human diabetes: hyperinsulinemia, mechanisms to increase insulin peptide bioavailability, and impaired insulin receptor sensitivity [4].

Enzymes that mediate oxidative stress are elevated in many longevity-conferring manipulations of *Drosophila* and *C. elegans* [18], and we see elevated *Sod-1* in both Types of long-lived *dInr* mutants. However, whether increased Sod-1 is sufficient to slow aging in *Drosophila* or mice is unresolved because its over-expression does not consistently extend lifespan [41]. Finally, fat body protein (*fbp*) was elevated in both types of *dInr*. Fat body protein was recently implicated as a longevity-limiting factor driven by juvenile hormone produced in young adults, where transiently blocking *Fbp* in young adults improves life expectancy [16]. Thus, increased *Fbp* in long-lived *dInr* genotypes is somewhat counter to expectation.

Few previously recognized aging-associated genes were seen to change only in the *dInr*^353^/*dInr*^wt^ adults. One exception involves the transcription factor *daughterless* (*da*): *da* expression was decreased and few binding sites for this transcription factor were identified among the differentially expressed genes of *dInr*^353^/*dInr*^wt^. Notably, depletion of the *daughterless* ortholog HLH-2 of *C. elegans* extends nematode lifespan [42].

There is a striking difference in innate immune related genes among the *dInr* longevity genotypes. In previous studies, the expression of antimicrobial peptides and peptidoglycan recognition proteins decline with age, while dietary and genetic manipulations that extend *Drosophila* longevity often maintain these defenses [43]. Here we see innate immune-associated genes are elevated in Type I *dInr*^E19^/*dInr*^74^ but not in Type II, and these expression differences correspond with the ability of adults to resist oral infection with *ECC15* bacteria. This outcome demonstrates that elevated innate immunity is not necessary for altered insulin signaling to extend lifespan.

Overall, while both Type I *dInr*^E19^/*dInr*^74^ and Type II *dInr*^353^/*dInr*^wt^ extend lifespan, these genotypes express sets of distinct and similar genes. Remarkably, their differences are caused by the placement of single amino acid substitutions within the insulin-like receptor.

### Methionine metabolism of Type I insulin-like receptor

We detected 170 metabolites from *dInr Drosophila* mutants assessed from a panel of 360 compounds. Sarcosine showed the most prominent difference: relative to wildtype, sarcosine was 7-fold higher in Type I adult soma. Sarcosine is produced when glycine is methylated by GNMT using S-adenosyl-methionine (SAM) as a substrate, and we found transcripts of *gnmt* were correspondingly elevated in *dInr*^E19^/*dInr*^74^. Serum sarcosine is normally reduced during mammalian aging, while it is increased in rodents when dietary restriction extends lifespan [44]. In cells and rats, sarcosine can induce autophagy and repress the activity of TOR [44]. In *Drosophila*, *gnmt* is regulated by the insulin-responsive transcription factor FOXO, and lifespan is increased when *gnmt* is over-expressed, while *gnmt* is required for a dominant-negative *dInr* transgene to extend lifespan [26].

Elevated sarcosine suggests the cells of *dInr*^E19^/*dInr*^74^ are processing excess methionine to keep SAM within homeostatic tolerances. SAM itself can potentially impact aging in several ways. Aside from the conversion of glycine into sarcosine, SAM is the key substrate for methylation of lipids, proteins and nucleotides, where such modifications alone could produce myriad impacts on aging [37]. SAM is also an entree molecule for the polyamine cycle when it is decarboxylated into an aminopropyl donor to produce spermidine and spermine. These polyamines are implicated to retard aging by modulating autophagy and lysosome biogenesis [45–47]. Finally, SAM in mammalian cells represses SAMTOR which otherwise will repress the activity of TOR [36]; the supply of SAM modulates the activation of mTORC1, which can thereby impact aging [36].

Donation of a methyl group by SAM produces SAH that is converted to homocysteine. Homocysteine is converted back to methionine or irreversibly directed into the transsulfuration pathway where its products may promote longevity by reducing ROS-associated damage [48]. In *Drosophila*, longevity conferred by dietary restriction is restored to wildtype level by inhibiting the rate-limiting transsulfuration pathway enzyme cystathionine β-synthase [49].

Aging itself alters the steady state levels of methionine cycle metabolites, as previously seen in *Drosophila* and in mammalian serum [50]. We find that Type I and wildtype adults have reduced somatic methionine and sarcosine at 30d relative to 15d. Yet, Type I adults retain transsulfuration product levels they expressed at younger age. These dynamics suggest that while aging reduces the absolute levels of methionine and SAM, cells in the older adults retain the ability to distribute carbons into the transsulfuration pathway.

We traced the fate of labeled carbons acquired from dietary methionine to verify that young Type I adults have elevated methionine cycle activity as suggested by their high level of sarcosine. When dietary methionine (labeled as m+5) traverses the central methionine cycle, it produces SAH with four labeled carbons that can be recycled to methionine, which is then labeled as m+4. The ratio of m+4 to m+5 (m+4/m+5) methionine therefore estimates the flow of carbons across the central methionine cycle [30]. Female *dInr*^E19^/*dInr*^74^ have strikingly greater m+4/m+5 relative to wildtype, indicating they metabolize a larger portion of methionine into SAH and subsequently back to methionine while also directing methionine towards the TSP.

### Methionine metabolism of the Type II (KID) insulin-like receptor genotype

Type II *dInr*^353^/*dInr*^wt^ affects the methionine cycle in remarkably different ways. The young adult soma has less methionine and sarcosine than wildtype, and transcripts for *gnmt* are not elevated. Young adults have less TSP products including methionine sulfoxide, cystathionine, cysteine, and taurine. In young adults there is less SAM than wildtype, while they have the same SAH as wildtype. The impact of age (30d) upon these metabolites is also distinct. Unlike wildtype and *dInr*^E19^/*dInr*^74^, the *dInr*^353^/*dInr*^wt^ adults retain their young (low) level of methionine but increase the amount of transsulfuration pathway products including cystathionine, methionine sulfoxide, cysteine, taurine, and hypotaurine.

We hypothesize these patterns might arise from the high level of egg production seen in *dInr*^353^/*dInr*^wt^. Young *dInr*^353^/*dInr*^wt^ females are 25% more fecund than wildtype [4] and this activity might sequester dietary methionine into eggs at a cost to somatic tissue. Old females produce fewer eggs, and dietary methionine may then flow into the TSP of somatic cells. In this model, the survival mechanisms supporting *dInr*^353^/*dInr*^wt^ longevity are biphasic: they first involve mechanism associated with low somatic methionine and later involve action derived from the TSP. The model contrasts with the results of Wei et al. [30] where dietary methionine restriction (MetR) elevated the m+4/m+5 ratio of methionine, whereas we find Type II decreases this ratio. This difference may arise because dietary restriction of methionine represses egg production, and paradoxically increase methionine cycle activity and the TSP, which is the very pattern we see for Type I *dInr*^E19^/*dInr*^74^. Type II, in contrast, is very fecund and this may reduce the ratio methionine m+4/m+5 ratio.

Isotope tracing of dietary methionine verifies these interpretations. Young adult *dInr*^353^/*dInr*^wt^ flies have a low m+4/m+5 methionine ratio, along with its low steady-state of TSP intermediates. These trends indicate there is either a relatively low activity of the methionine cycle beyond SAM or, SAM is flowing into the polyamine pathway. Our data suggest that *dInr*^353^/*dInr*^wt^ directs SAM to the polyamine pathway because we find elevated m+1 MTOB and m+1 MTA in both young and old Type II adults. Although the proportion of M+5 methionine increases in old (30d) Type II adults, we could not calculate m+4/m+5 because the level of m+4 methionine is negligible. These data suggest that carbons from dietary methionine are shunted into the salvage pathway (indicated by increased m+1 labeling in MTA, MTOB, and SAH), which can be a source of polyamines. Considering the reported pro-longevity effects of spermidine [46, 47, 51], our observations may partially explain how *dInr*^353^/*dInr*^wt^ is long-lived. Thus, young *dInr*^353^/*dInr*^wt^ experience restricted somatic methionine that impedes activity of the central methionine cycle and they increase the flow methionine into the salvage pathway. Products of the TSP increase as these flies age, which may foster their response to stress.

### Methionine restriction and aging

Dietary methionine restriction (MetR) slows aging across many species including rodents and *Drosophila* [52–55], as does direct genetic manipulation of the methionine cycle. Knockdown of *C. elegans* S-adenosyl methionine synthetase (*sams-1)* increases adult lifespan [56]. *Drosophila* lifespan is increased 25% by expressing an exogenous methioninase that degrades methionine [29]. MetR decreases levels of intermediates in the methionine metabolism pathway including methionine, methionine sulfoxide, SAM, SAH, cystathionine, and 5’-methylthioadenosine (MTA) [29, 30, 57]. Profiles of methionine metabolism are also altered in long-lived species and strains, such as naked mole rats [58], flies selected for delayed reproductive senescence [28, 59], and Ames mice [60]. Across these cases, methionine restriction is thought to slow aging through many potential avenues, including protein and nucleotide methylation, polyamine production, protein translation, and altered energy or oxidative status [50].

Steady-state metabolomics and tracing analysis demonstrate that young Type II adults have attributes of methionine metabolism that are associated with extended lifespan. The long-lived *dInr*^353^/*dInr*^wt^ adults have less steady-state methionine (MetR-like), less steady-state methionine sulfoxide, increased m+1 labeling in the salvage pathway intermediates and in SAH, and elevated age-associated transuslfuration products.

We conducted genetic epistasis analysis to verify that *dInr*^353^/*dInr*^wt^ extends longevity because it modifies the central methionine cycle activity. Unlike where *gnmt* has been reported as required for repressed insulin signaling to slow aging [25, 26], the extended longevity of *dInr*^353^/*dInr*^wt^ was independent of *gnmt*. As well, Type II *dInr*^353^/*dInr*^wt^ did not interact with SAMTOR or S6K to slow aging, suggesting its impact is independent of TOR. Rather, *dInr*^353^/*dInr*^wt^ requires *ahcy13*. AHCY-13 encodes SAHH (S-adenosylhomocysteine hydrolase), which converts SAH into homocysteine. Emerging work shows that AHCY itself modulates aging: *Drosophila* lifespan was increased by downregulating genetic repressors of AHCY [28], and *C. elegans* lifespan was increased by a deficiency of AHCY-1 that increases the ratio of SAH to SAM [61]. Cells normally maintain low concentrations of SAH because SAH is an inhibitor of SAM-dependent methylation reactions. Inhibition of AHCY blocks SAH clearance and prevents the flux of methionine through the methionine cycle and its subsequent flux into the transsulfuration pathway. We have previously demonstrated that downregulation of Ahcy13 in *Drosophila* dramatically upregulates SAH levels [28], thus strongly inhibiting methionine flux. By inhibiting Ahcy13 in wildtype and *dInr*^353^ flies, we asked if the observed changes in methionine metabolism are required for the lifespan extension in *dInr*^353^/*dInr*^wt^. Depleting *ahcy13* shortens *dInr*^353^/*dInr*^wt^ lifespan to the level of similarly manipulated wildtype controls. We conclude that *dInr*^353^/*dInr*^wt^ extends longevity, in part, because it reprograms activity of the methionine cycle. This reprogramming has three impacts: i) methionine is routed towards the salvage/polyamine pathway; ii) the flux of the central via methionine cycle is reduced, and iii) old adults preserve an ability to upregulate production of intermediates in the transsulfuration pathway.

### Alternative modes of longevity control by insulin/IGF signaling

Canonical Type I insulin mutants persistently elevate the products of the TSP relative to wildtype. These mutants have reduced fecundity, perhaps sparing the allocation of dietary methionine to eggs and directing it to somatic TSP. The *dInr*^E19^/*dInr*^74^ mutant continuously elevates cysteine, taurine, and reduced-glutathione, which may favor somatic maintenance by managing the damage of ROS [48] [62] [63] [64]. The Type II mutant at the Kinase Insert Domain is substantially different. The females are very fecund, and young adults have reduced somatic methionine activity and less TSP. Yet, the mortality rate at these ages is remarkably low [4], which may arise from the longevity assurance mechanisms of dietary methionine restriction, polyamine synthesis, and decreased methionine sulfoxide. Furthermore, the metabolism of these flies is flexible. As they age, Type II adults increase their (relative) production of TSP, perhaps because reduced egg production with age frees methionine for somatic tissue. The different metabolic syndromes among the Type I and Type II *dInr* arise from a change in the placement of a single amino acid within the receptor. This structure-to-function mapping suggests the kinase insert domain regulates aging without insulin-resistance because it reduces somatic methionine cycle activity in young adults, in contrast to the excess methionine and high activity seen in canonical longevity alleles of *dInr*.

The ability to efficiently adapt metabolism by substrate utilization in response to nutrient intake and timing is known as metabolic flexibility, as first describe in lean humans as they adapt their metabolic fuel preference during fasting and insulin infusion relative to the static responses of obese individuals [65, 66]. Metabolic flexibility may partially explain the paradox that mild insulin resistance has been observed in a variety of long-lived mice and is therefore not necessarily an indicator of poor health or shortened lifespan [67]. Metabolic flexibility is typically applied to the scales of daily feeding and circadian rhythms, where tissues transit from carbohydrate to lipid metabolism. Here, we describe metabolic flexibility from an insulin sensitive mutant in the dInR Kinase Insert Domain on the scale of aging. This flexibility permits adults to vary their methionine metabolism from a MetR-like state when young to metabolism where methionine flux is increased into the transsulfuration pathway.

## Materials and Methods

### Fly stocks and culture

The insulin receptor mutants were generated and described in [4]: *dInr*^+[HR]^ (wildtype, accession ‘29B’), *dInr*^353[HR]^, *dInr*^E19[HR]^, *dInr*^74[HR]^. All alleles were generated by ends-out gene replacement. Wildtype accession ‘29B’ is a substitution by gene replacement of the wildtype allele back into the originating stock (*w*Dahomey, *w*Dah, source: L. Partridge UCL, UK). Mutant alleles were produced by nucleotide substitutions of the *w*Dah wildtype allele to alter the targeted amino acid. Mutant *dInr* alleles were maintained as third chromosome (TM6) balanced stocks. Mutant genotypes for all analyses were generated by crossing *dInr*^353^/TM6B *Tb,Sb,e* to *dInr*^wt^, recovering F1 *dInr*^353^/*dInr*^wt^, and by crossing *dInr*^E19^/*Tb,Sb,e* to *dInr*^74^/*Tb,Sb,e*, recovering trans-heterozygote *dInr*^E19^/*dInr*^74^. Stocks used in the epistasis analysis: *gnmt*-RNAi: w[1118]; P{GD10563}v25983 (VDRC); *samtor*-RNAi: P{KK107414}VIE-260B (VDRC); Ahcy13-RNAi: *y*^1^ *sc*\* *v*^1^ *sev*^21^; P{TRiP.HMS05799}attP40 (BDSC #67848); S6K: w[1118]; P{w[+mC]=UAS-S6k.TE}2 (BDSC #6912). Each of the VDRC and BDSC lines were backcrossed by *w*Dah;+/+;+/+ for 5 generations. Flies were reared at 25°C, 40% RH, 12L:12D on standard food media: cornmeal (5.2%), sugar (11.0%), autolyzed yeast (2.5%; SAF brand, Lesaffre Yeast Corp., Milwaukee, WI, USA.), agar (0.79%) (w/v in 100 mL water) with 0.2% Tegosept (methyl4-hydroxybenzoate, Sigma, St Louis, MO, USA), with baker’s yeast supplemented to the food surface.

### Infection survival assay

We followed the protocols of Troha and Buchon [68] to orally infect adults with *Pectinobacterium carotovora carotovora* (strain ECC15). ECC15 was grown in shaking culture at 29C overnight, pelleted by centrifugation (10 min at 2400g, 4C) and resuspended in sterile PBS to OD600. Treatment conditions were initiated with 8-to-9-day-old adult females, collected over a period of 48 hours post-eclosion in the presence of males. The beginning of the treatment period is designated as survival age zero. Cohorts of 100 adults for each genotype were evenly distributed across 10 food vials per treatment group (PBS control, ECC15). A volume of 100 uL of either sterile PBS or Ecc15 suspension was added to a round filter paper placed on top of the food within each vial. Prior to the treatment, flies were starved for 2 hours in empty vials, then placed in ECC15 or control treated vials and retained for 18-24 hours. Subsequently, adults were flipped into fresh, uninfected food daily, at which time dead flies were removed and counted. We plotted Kaplan-Meier survival curves based on right-censored data at survival age 47 days. Cox-proportional survival analysis was used to evaluate genotype, treatment and genotype-by-treatment interaction.

### RNA-seq data and analysis

Flies eclosed and mated over a 48-hour period and females of each genotype (wildtype, *dInr*^E19^/*dInr*^74^, *dInr*^353^/*dInr*^wt^) were the collected into six biological replicates of 20 females. Each replicate was lysed in Trizol using a bead Tissuelyzer (Qiagen) at room temperature. RNA was extracted from each sample by NH_4_OAc and EtOH precipitation. RNA concentration (ng/ul) and purity (absorbance: 260nm/280nm) was confirmed with a Nanodrop spectrophotometer (ThermoFisher Scientific). Samples were sent to Genewiz for library preparation (Illumina, RNA with PolyA selection) and Illumina HiSeq 2×150 bp sequencing. Returned reads were processed on the Basepair platform for RNA-Seq QC, alignment, and counts. Differential expression, gene set enrichment, K-means enrichment and transcription factor binding site enrichment were generated with iDEP (http://bioinformatics.sdstate.edu/idep96/).

### Steady State Metabolomics

Tissue for analysis of steady state metabolomic profiles were collected from females at age 15d and 30d. Females eclosed in the presence of males and were flipped into female only vials from age 2 days until sample collection. Six biological replicates of each genotype (*dInr*^wt^/*dInr*^wt^, *dInr*^E19^/*dInr*^74^, *dInr*^353^/*dInr*^wt^) were generated with 10 females per sample. At age 15 and 30d, females were collected between 9 and 10 AM, anesthetized with CO_2_ and dissected to remove the abdomen to avoid signal from eggs. The somatic tissue was flash frozen in liquid nitrogen and stored at -80. Samples were shipped on dry ice to the Northwestern Metabolomics Research Center (NWMRC) at the University of Washington, Seattle. Protein from each biological sample was extracted at the NWMRC and quantified by Targeted LC-MS across a 300-feature set (https://northwestmetabolomicsorg.wpcomstaging.com/300-aqueous-metabolites/). The analysis returns Quality Control statistics and relative concentration of each detected metabolite based on isotope-labeled internal standards.

Data processing and analyses were performed using metaboanalyst (http://www.metaboanalyst.ca/MetaboAnalyst/). Metabolites that were not detected in 40% of the samples were excluded from the analysis. Data were normalized to the median (per sample) and processed through log transformation. Heat map and hierarchical clustering was generated using Pearson correlations and Ward’s method. Metabolite Set Enrichment Analysis (MSEA) was performed using Metaboanalyst which uses the KEGG (http://www.genome.jp/kegg/pathway.html) pathway database. Metabolite sets containing at least 5 compounds were employed in the analysis. MSEA calculates hypergeometric test scores based on cumulative binominal distribution.

### Metabolite profiling labeled samples

Samples were mixed with −20 °C 80%:20% methanol:water (extraction solvent), vortexed, and immediately centrifuged at 16,000 x g for 20 min at 4°C. The supernatant was collected for LC-MS analysis. LC was performed on an Xbridge BEH amide HILIC column (Waters) with a Vanquish UHPLC system (Thermo Fisher). Solvent A was 95:5 water: acetonitrile with 20 mM ammonium acetate and 20 mM ammonium hydroxide at pH 9.4. Solvent B was acetonitrile. The gradient used for metabolite separation was 0 min, 90% B; 2 min, 90% B; 3 min, 75%; 7 min, 75% B; 8 min, 70% B, 9 min, 70% B; 10 min, 50% B; 12 min, 50% B; 13 min, 25% B; 14 min, 25% B; 16 min, 0% B, 21 min, 0% B; 21 min, 90% B; and 25 min, 90% B. MS analysis was performed on a Orbitrap Exploris 480 mass spectrometer (Thermo Fisher) by electrospray ionization with parameters as: scan mode, full MS; spray voltage, 3.6 kV (positive) and −3.2 kV (negative); capillary temperature, 320 °C; sheath gas, 40 arb; aux gas, 7 arb; resolution, 120,000 (full MS); scan *m*/*z* range, 70 to 1,000.

Raw LC–MS data files were first converted to mzXML format files by ProteoWizard. Then, the data were analyzed using El-MAVEN Software (Elucidata; elucidata.io). For tracer experiments, isotope labeling was corrected for the natural abundance of ^13^C isotopes.

### Demographic epistasis analysis

Females for survival analysis were collected as F1 progeny over a 48-hour emergence period in the presence of males, then separated into all female demography cages with 125 adults in each of three cages per genotype. Demography cages were made from 1 L clear food service containers with a ventilated lid, a gasket covered aperture to provide access to remove dead flies and a port near the cage bottom that opened to a plastic tube affixed a standard glass media vial with 3 ml of standard Drosophila diet. Each two days, dead flies were removed by aspiration and counted, and fresh food vials were provided.

Genotypes for survival analyses were produced by replacing the balanced third chromosome of *dInr*^353^ with the *tubulin*-Gal4.Gal80 third chromosome, followed by replacing the second chromosome of offspring with the targeted UAS-RNAi or over-expression. Females from these crosses were set up in controlled density bottles to lay eggs. Hatching larvae were reared at 25°C to suppress transgene expression. Emerging adults were shifted to the permissive temperature (29°C) and sorted into demography cages, maintained at 29°C. Wildtype F1 control and *dInr*^353^ single mutant genotypes were generated by replacing the third chromosome balancer with the *tubulin*-Gal4,Gal80 third chromosome.

## Acknowledgments

This work was supported by NIGMS R35 GM146869 (A.P.), NIA R01 AG082801 (A.P. and M.T.), NIA R03 AG075651 (A.P.), R03 CA286521 (A.P.), Richard King Mellon Foundation award (A.P.), NAM Healthy Longevity Catalyst Award (A.P.), NIA R01 AG059563 (M.T.), NIA R01 AG069639 (M.T.).

## Data and materials availability

All data are available within the Article and Supplementary Files, or available from the authors upon a reasonable request.

## Figure Legends

**Supplementary Figure 1.**
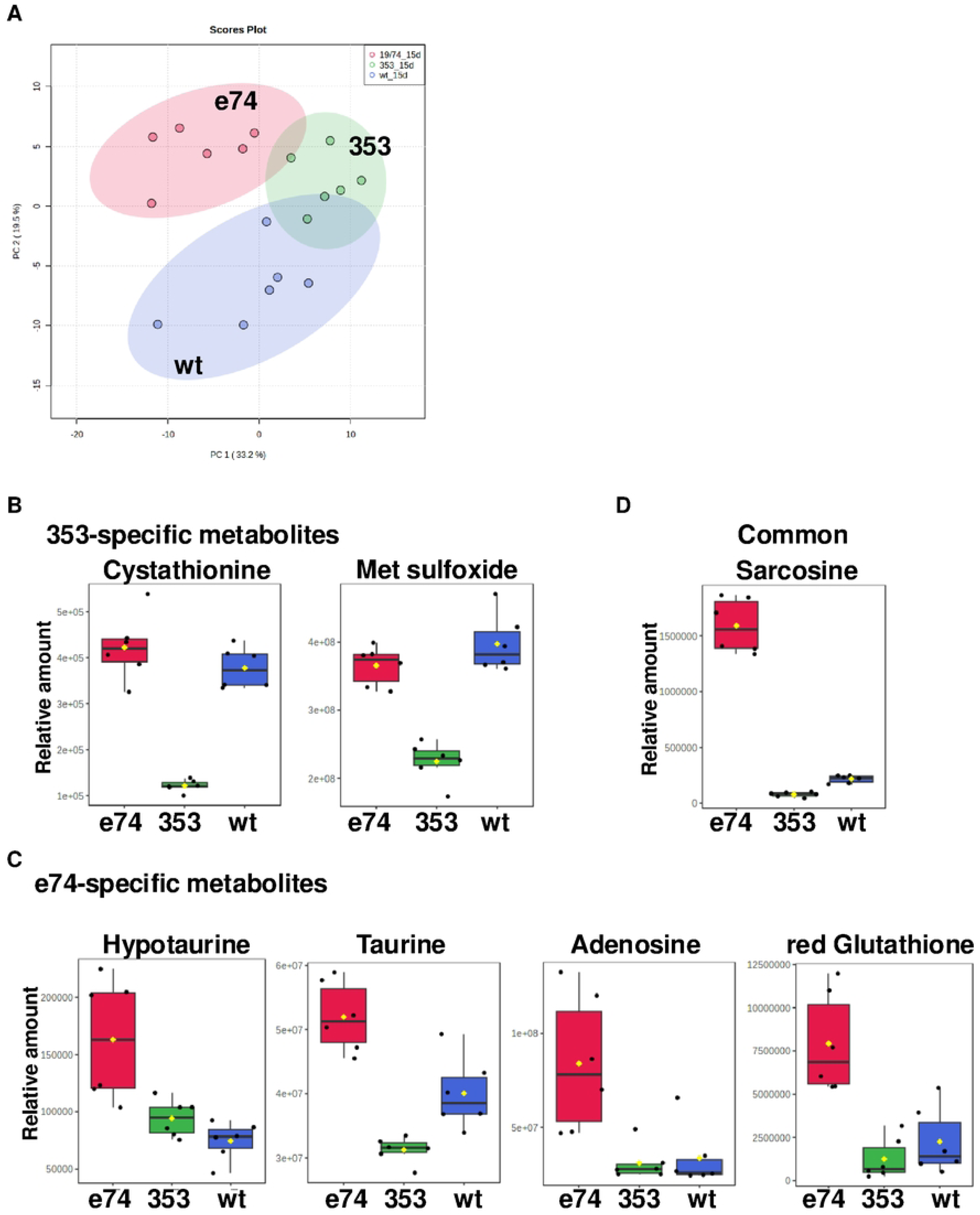
Metabolomic profiles of dInr^E19^/dInr^74^ and dInr^353^/dInr^wt^. (**A**) Principal component analysis of metabolomic profiling in *Type I dInr^E19^/dInr^74^ and Type II dInr*^353^/*dInr*^wt^ genotypes. (**B**) Relative levels of metabolites (cystathionine and methionine sulfoxide) significantly altered in *dInr*^353^/*dInr*^wt^ (Type II) genotype. (**C**) Relative levels of metabolites (hypotaurine, taurine, adenosine, reduced glutathione) significantly altered in *dInr^E19^/dInr^74^* (Type I) genotype. (**D**) Relative levels of metabolites (sarcosine) significantly altered in both *Type I dInr^E19^/dInr^74^ and Type II dInr*^353^/*dInr*^wt^ genotypes.

**Supplemental Table 1. Differentially expressed genes**.

List of differentially expressed genes in *Type I dInr^E19^/dInr^74^ and Type II dInr*^353^/*dInr*^wt^ genotypes.

**Supplemental Table 2. Gene set enrichment analysis**.

Gene set enrichment analysis in *Type I dInr^E19^/dInr^74^ and Type II dInr*^353^/*dInr*^wt^ genotypes.

**Supplemental Table 3. List of significantly altered metabolites**.

List of significantly altered metabolites in *Type I dInr^E19^/dInr^74^ and Type II dInr*^353^/*dInr*^wt^ genotypes.

